# Facile discovery of isonitrile natural products *via* tetrazine based click reactions

**DOI:** 10.1101/711853

**Authors:** Yaobing Huang, Wenlong Cai, Antonio Del Rio Flores, Frederick Twigg, Wenjun Zhang

## Abstract

A facile method for the quick discovery and quantification of isonitrile compounds from microbial cultures was established based on the isonitrile-tetrazine click reaction. A in situ reduction further enabled bioorthogonal ligation of primary and secondary isonitriles for the first time.

---

Isonitrile-containing natural products have exhibited intriguing bio-activities ranging from antibiotics, metal acquisition, detoxification and virulence.^[1,2]^ Up to date, over 200 isonitrile-containing natural products have been identified,^[3]^ including a well-known antiviral agent xanthocillin from *Penicillium notatum*,^[4]^ and epoxy isonitrile compounds such as aerocyanidin, YM-47515 and amycomicin which displayed potent antibiotic activities against Gram-Positive pathogens **(Figure 1)**.^[5-7]^ Recently we have discovered a family of isonitrile lipopeptides (INLPs) synthesized from five conserved enzymes widely spread in actinobacteria, which have been indicated to be related to virulence in pathogenic mycobacteria.^[8]^ A similar INLP, SF2768, was also reported recently to function as a chalkophore that mediates copper acquisition in *Streptomyces thioluteus*.^[9]^

**Figure 1.**
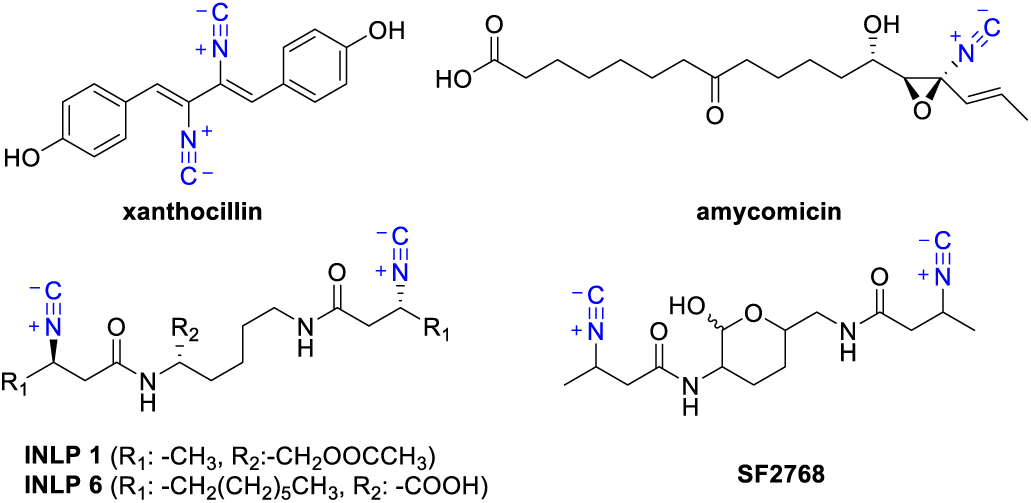
Scaffolds of isonitrile natural products.

We are interested in the discovery of isonitrile-containing natural products because of their high potential as pharmaceutical agents as well as building blocks in synthetic chemistry.^[10,11]^ Currently, there are two biosynthetic mechanisms known to generate an isonitrile functionality in nature: the isonitrile synthases (represented by IsnA),^[12,13]^ and the non-heme iron(II) dependent oxidases/decarboxylases (represented by ScoE)^[14]^. Microbes continue to serve as an important source for future discovery of isonitrile natural products as genomic analysis suggested that these putative isonitrile biosynthetic genes are widely spread but the associated metabolites are often unknown.

The isonitrile moiety is labile and has weak UV-Vis signatures, making purification and structural elucidation of isonitrile natural products quite challenging. We then aim to develop a facile and selective method for isonitrile detection based on its unique reactivity. Tetrazines have been widely used in click reactions with strained alkenes/alkynes to allow efficient bioorthogonal ligation for protein labelling.^[15-17]^ Similarly, it has been adopted in a [4+1] cycloaddition click reaction with isonitrile for the bioorthogonal ligation of a tertiary isonitrile-tagged protein.^[18-20]^ It’s worth noting that spontaneous hydrolysis of the tetrazine-isonitrile conjugate was observed to release an aminopyrazole motif when a primary or secondary isonitrile compound was used, which restricted a broad application of this conjugation reaction.^[18, 21]^ Nonetheless, we reason that this reaction could be adapted for isonitrile natural product discovery from microbial cultures due to the following characteristics: 1) It has been demonstrated to be a rapid reaction under mild conditions. 2) The reaction may be associated with a color change for tetrazine,^[18,22]^ which could serve as an early indication of the presence of isonitrile. 3) While release of an aminopyrazole product is not ideal for a bioorthogonal ligation, this specific product delivers a good diagnostic tool to confirm the presence of an isonitrile compound (**Figure 2**). We here thus aim to optimize the isonitrile-tetrazine click reaction to enable the facile detection of isonitrile metabolites produced by microbes.

**Figure 2.**
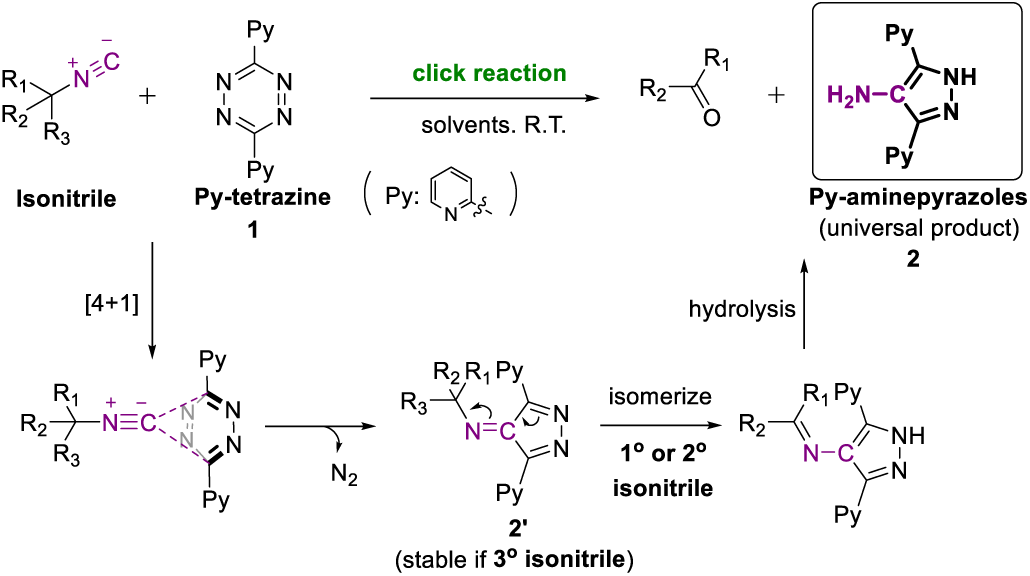
Click reactions between tetrazine and different isonitriles.

Previous studies on the Py-tetrazine (**1**) and isonitrile reaction suggested that the reaction rates were affected by both the solvents and isonitrile substrates, motivating a thorough kinetic investigation.^[17,18]^ The formation of compound **2** was monitored and quantified using LC-HRMS analysis (**Figure S1**). Reactions in several commonly used analytical reagent (AR) grade organic solvents using p-toluenesulfonylmethyl isocyanide (**3)**, a primary isonitrile compound, showed different kinetic curves, with much higher rates in dimethylsulfoxide (DMSO) and methanol (MeOH), and lower rates in tetrahydrofuran (THF) and acetonitrile (MeCN) (**Figure 3 and Table 1**). These data implied a beneficial effect of polar aprotic solvents on the click reaction. By employing water as the co-solvent to these organic solvents (1:1, v/v), enhanced reaction rates were observed (**Table 1**). These results confirmed that polar aprotic solvents promote the reaction rate of the tetrazine-isonitrile reaction, supporting the application of the tetrazine probe directly in microbial cultures (water-based) for isonitrile detection. It is notable that **2** disappeared after a prolonged incubation in DMSO, likely due to oxidation. Slow reaction rates were observed for **3**, which was likely due to the formation of an α, β-unsaturated conjugate that slows down the subsequent hydrolysis (**Figure 3)**. This was confirmed by using an alternative isonitrile standard, cyclohexyl isocyanide (**4**), which showed significantly higher rate constants than **3** under the same reaction conditions (**Table 1**). Despite the nature of the isonitrile substrate, full conversion of both substrates (1 mM) by 10 mM Py-tetrazine was observed in MeOH-H_2_O (1:1, v/v) within one hour, suggesting that this reaction condition could be used to quantify the amount of isonitrile metabolites from cell cultures for a concentration of up to 1 mM. It is notable that tetrazine dissolved in a variety of solvents showed a bright-red color, which quickly changed into light yellow when the click reaction proceeded to form **2** (light yellow) (**Figure S2)**, potentially useful for quick screening of the presence of isonitrile-containing compounds.

**Table 1.**
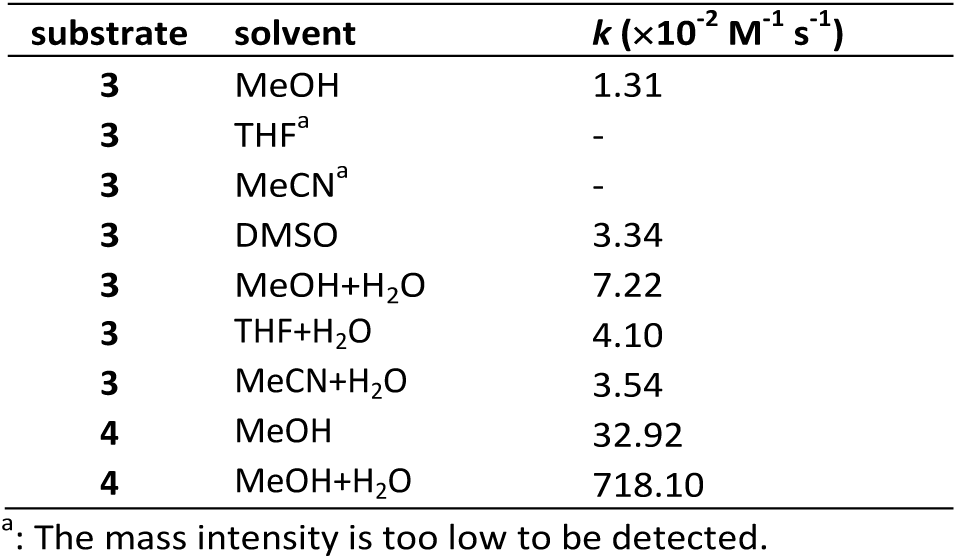
Rate constants of click reaction in different solvents.

**Figure 3.**
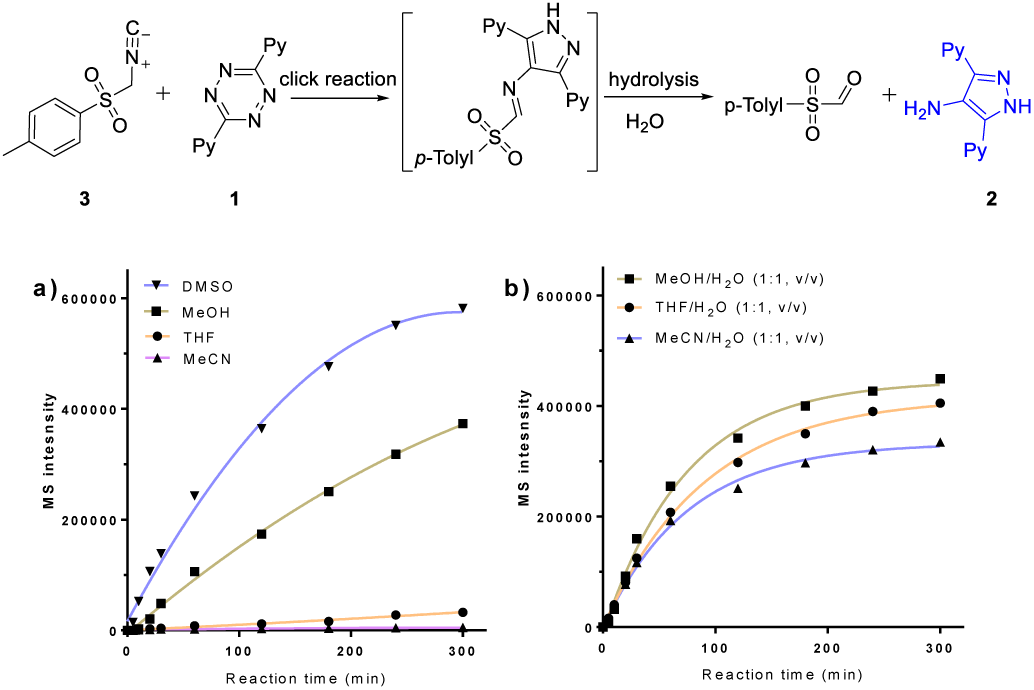
Solvent effect of the isonitrile-tetrazine click reaction. a) reaction profiles in different organic solvents. b) reaction profiles in different mixed solvents. Condition: isonitrile **3** 5 mM, Py-tetrazine 10 mM, solvent (AR grade), R.T.

We next tested the reaction for isonitrile natural product detection and quantification from microbial cultures using two of our previously developed INLP production strains, *E. coli-ScoA-E* and *E. coli-MmaA-E*, which produce INLP 1 and INLP 6 (**Figure 1**), respectively.^[8]^ After adding Py-tetrazine to the culture extracts, a quick color change of the reaction solution from red to pink was firstly observed with bare eyes, which was not detected when culture extracts of the control strain of *E. coli* harboring blank vectors were used. The LC-HRMS analysis of reaction mixtures revealed the production of the universal product **2** from *E. coli-ScoA-E* and *E. coli-MmaA-E*, but not from the *E. coli* control strain that does not produce isonitrile-containing metabolites (**Figure S3, S4**), demonstrating the selectivity of this reaction toward isonitrile groups. Neither INLP 1 nor INLP 6 were detected after reacting with Py-tetrazine (10 mM) (**Figure S3, S4**), suggesting that a complete conversion of both isonitrile metabolites were achieved. Subsequent determination of the amount of generated **2** using a calibration curve (**Figure S1)** led to calculated production titers of INLP 1 and INLP 6 to be 4.6±0.3 and 39.3± 1.0 µM, respectively (**Figure S5**).

We then applied this reaction to unknown isonitrile natural product discovery through genome mining. We targeted *Streptomyces tsukubenesis NRRL 18488* as its sequenced genome contains a gene cluster that is predicted to produce an INLP.^[9]^ This strain is well-known for producing the clinically important immunosuppressant tacrolimus (FK506),^[23-25]^ as well as the acyl-CoA synthetase inhibitors triacsins,^[26]^ but no INLP has been reported to be produced by this strain. *S. tsukubenesis* was cultured in six different media which were then subjected to the tetrazine click reaction. Two (ISP2 and YD) out of the six media displayed an obvious color change, suggesting the presence of an isonitrile metabolite. The subsequent LC-HRMS analysis confirmed that compound **2** was generated in the reactions with culture extracts grown in ISP2 and YD, but not detected with culture extracts grown in the other four media (**Figure S6**), consistent with the observed color change. To reveal the identity of the produced INLP, we performed a comparative metabolomic analysis of the culture extracts before and after the tetrazine reaction, and identified two metabolites with the same mass ([M+H^+^] 337.1882) that were present before the reaction but completely disappeared after (**Figure 4 and S7**). Database search suggested that these two metabolites are two known SF2768 anomers (**Figure 1**). SF2768 was firstly reported in 2011 as a cryptic antibiotic ^[27]^ and recently linked to an INLP gene cluster in *S. thioluteus*.^[9]^ The known biosynthetic genes required for its biosynthesis were found to be conserved in the genome of *S. tsukubenesis*, consistent with the production of SF2768 also in *S. tsukubenesis* (**Figure 4e**). We further determined the titers of this metabolite to be ∼1.6 µM in both producing media (**Figure S5**). We thus demonstrate that this click reaction could serve as a convenient screening method for culturing conditions for isonitrile production, as well as facilitating the isonitrile metabolite identification by comparative metabolomics.

**Figure 4.**
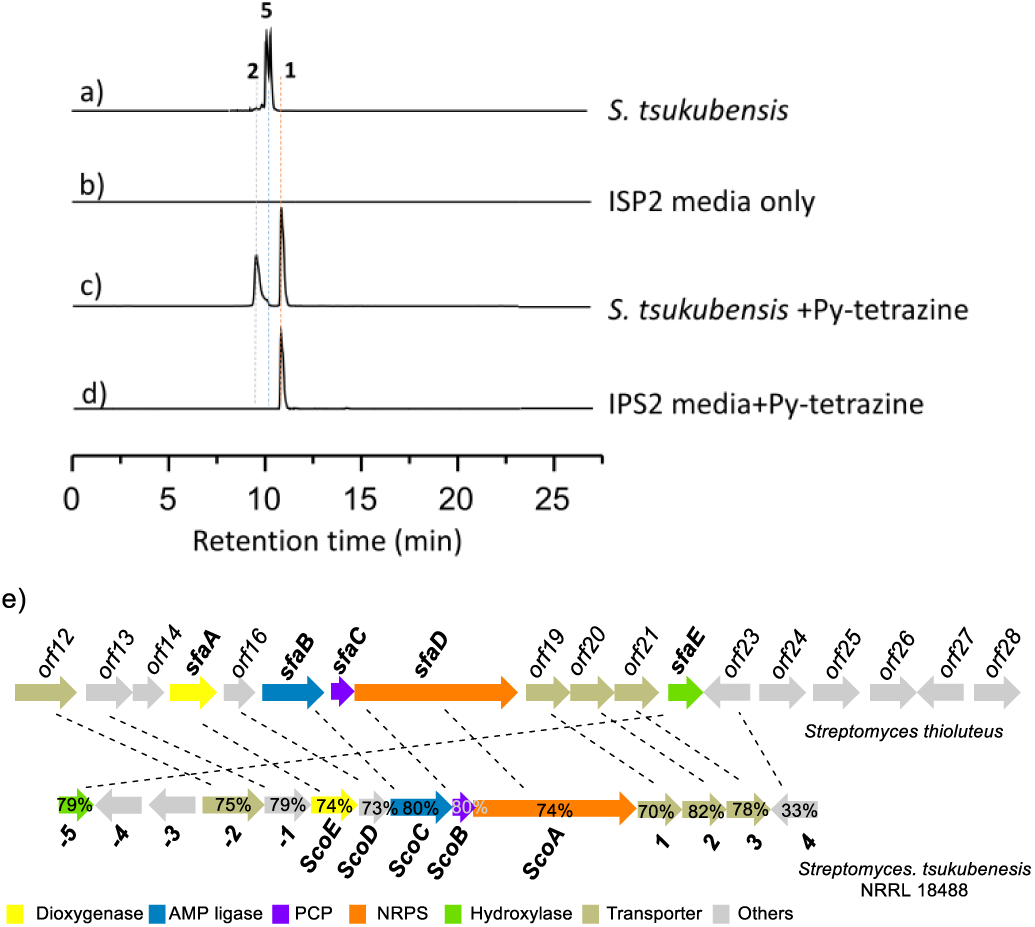
Identification of isonitrile compound from *Streptomyces tsukubenesis*. Extracted ion chromatograms showing a) the presence of isonitrile compound SF2768 in *S. tsukubenesis*, b) no target compound in ISP2 media only, c) full isonitrile conversion with the production of universal product **2** after the addition of Py-tetrazine, and d) no reaction if Py-tetrazine was added the ISP2 media. e) Homologous genes between the two clusters are cross-linked and the similarities were provided. The calculated masses for SF2768 (337.1870), Py-tetrazine (237.0883), and universal product (238.1087) with 20 ppm mass error tolerance were used for each trace.

While the production of the universal aminopyrazole product is convenient for the indication of the presence of primary and secondary isonitrile metabolites, bioorthogonal ligation without hydrolysis would offer additional benefits, such as to directly capture and enrich isonitrile metabolites from crude extracts and to facilitate the conjugate detection and purification via a unique chromophore. ^[18-20]^ We then tried to modify the isonitrile-tetrazine reaction to maintain the conjugate product and minimize the subsequent hydrolysis. To achieve this goal, we selected NaBH_3_CN, a mild reductant that was known to be able to selectively but slowly reduce imines^[28]^. We reasoned that to favor reduction over hydrolysis, a solvent in which the formation rate of **2** is low would be preferred. We thus selected MeCN as the solvent since it showed the lowest formation rate of **2** based on the previous reactions with different solvents (**Table 1**). *E. coli-MmA-E* culture extracts containing INLP 6 were then subjected to the tetrazine click reaction in MeCN, with the addition of NaBH_3_CN (**Figure 5**). LC-HRMS analysis of the reaction mixture revealed that an expected reduced conjugate (**6**) was successfully formed (calculated: [M+H^+^] 715.4654, found: 715.4635) (**Figure 5 and S8**). Further quantification revealed that approximately two thirds of conjugates were reduced by NaBH_3_CN (30 mM) in the presence of 10 mM Py-tetrazine. Similar results were observed in the *in-situ* reduction of the click reaction between Py-tetrazine and INLP extracts from *E. coli-ScoA-E* and *S. tsukubenesis*, respectively (**Figure S9, S10**). We have thus demonstrated an effective method to capture secondary isonitrile metabolites which broadens the scope of the tetrazine-isonitrile click reaction for bioorthogonal ligation applications.

**Figure 5.**
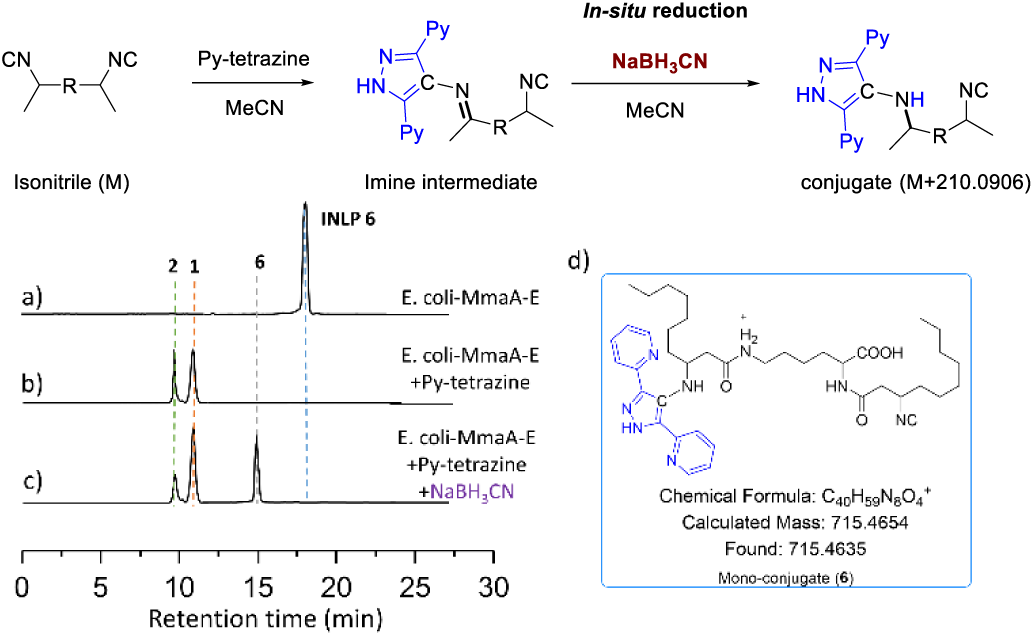
*In-situ* reduction of the click reaction between *E. coli-MmaA-E* extracts and Py-tetrazine with NaBH_3_CN. a) Extracted ion chromatograms showing the production of INLP 6 in *E. coli-MmaA-E* extracts. b) Extracted ion chromatograms showing the full conversion of INLP 6 and the presence of the universal product **2** after the click reaction of *E. coli-MmaA-E* extracts with Py-tetrazine. c) Extracted ion chromatograms showing the production of mono-conjugate **6** after the addition of NaBH_3_CN to the click reaction. The calculated masses for INLP 6 (505.3748), Py-tetrazine (237.0883), universal product (238.1087) and mono-conjugate **6** (715.4654) with 20 ppm mass error tolerance were used for each trace. d) a putative molecular structure of mono-conjugate **6**.

## Conclusions

In summary, we have reported a facile method for the discovery of isonitrile-containing natural compounds produced by microbes based on the isonitrile-tetrazine click reaction. The convenient colorimetric assay serves as a quick indication of the presence of isonitrile compound from microbial cultures, and the production of a universal aminopyrazole product confirms the presence of the primary or secondary isonitrile metabolites. We also demonstrated that a complete conversion of isonitrile metabolite can be readily achieved in culture extracts to allow a quantitative readout of the amount of isonitrile compounds. We further extend this reaction to include an *in-situ* reduction of conjugates, thus gaining access to the bioorthogonal ligation for different types of isonitrile compounds beyond the current limitation of tertiary isonitrile compounds only. This study thus paves the way for future discovery and characterization of isonitrile-containing compounds which have broad medical and biotechnological applications.

## Conflicts of interest

There are no conflicts to declare.

## Acknowledgement

This research was financially supported by grants to Y.H. from the QinLan project (Jiangsu Province), and to W.Z. from the NIH (DP2AT009148), Alfred P. Sloan Foundation, and the Chan Zuckerberg Biohub Investigator Program.

