## Supplementary materials for "Facile discovery of isonitrile natural products *via* tetrazine based click reactions"

### Supporting information

I. Materials

II. Methods

III. Standard curves for HPLC analysis.

IV. Click reaction of INPLs from *E. coli-ScoA-E* and *E. coli-MmaA-E*.

V. Identification of isonitrile compounds from *Streptomyces tsukubanesis NRRL 18488.*

VI. *In-situ* reduction of the click reaction.

VII. Titer quantification of isonitrile compounds from strain culture extracts.

#### I. Materials

3,6-Di-2-2pyridyl-1,2,4,5-tetrazine (Alfa Aesar, 96%), 4-toluenesulfonylmethylisocyanide (Oakwood Chemical, 98%), cyclohexyl isocyanide (Sigma, 98%), all the other solvents (A. R grade) and reagents were bought from local chemical companies and used without further purification.

Media components used in the paper were listed.

| Media | Composition (g/L) |
|---|---|
| A | Glucose (10), yeast extract (5), starch (20), peptone (5), NaCl (4), $K_2HPO_4$ (0.5) $MgSO_4$ (0.5), $CaCO_3$ (2) |
| ISP2 | Yeast extract (4), malt extract (10), glucose (4) |
| J | Tryptone (5), malt extract (3) glucose (11), yeast (3) |
| R5 | Sucrose (103), $K_2SO_4$ (0.25), $MgCl_2$ (10.1), Glucose (10), Casaminoacids (0.1), Trace element solution (2 ml), yeast extract (5), TES buffer (5.73) |
| TSB | Commercially available |
| YD | yeast extract (5), maltose (10), glucose (4), $MgCl_2$ (2), $CaCl_2$ (1.5) |

#### II. Methods

**Click reaction between isonitrile and tetrazine:** Typically, 10 mM isonitrile compound and 20 mM tetrazine in specific solvent were prepared and mixed in equivalent volume. The mixture was stirred at room temperature for a certain reaction time and sampled for Agilent 6520 HPLC-MS Q-TOF or SingleQ analysis.

The universal product was separated using an Agilent 1260 HPLC with a C18 Vydac 218TP1022 column 10 μm (22 x 250mm) using a linear gradient of 2-98% $CH_3CN$ (vol/vol) over 20 min in $H_2O$ with 0.1% formic acid at a flow rate of 3 mL/min. The product was a light yellow solid.

**Culture condition for production of INLPs**: *E. coli-ScoA-E* and *E. coli-MmaA-E* were grown triplicate in LB media, respectively. Typically, 30 mL cultures were grown at 37 $^{o}$C in LB containing appropriate antibiotics (for *E. coli-ScoA-E*, Kar, Carb, Spec and Cm; for *E. coli-MmaA-E*, Kar, Carb, Spec) until reaching the $OD_{600}$ value 0.5. Then, for *E. coli-ScoA-E*, 2, 3-trans-crotonic acid, Glycine and IPTG were added to the culture with a final concentration of 1, 10 and 0.5mM for induction, respectively; The same procedure was applied to *E. coli-MmaA-E* except decenoic acid was used instead of 2,3- trans-crotonic acid. After induction, the temperature was decreased to 20 $^{o}$C and the culture was shaken for another 48 h to allow the production of compounds. The culture was extracted by two equal amounts of ethyl acetate, dried and redissolved in methanol for LC-HRMS analysis. LC-HRMS was conducted on an Agilent Technologies 6520 accurate-mass Q-TOF instrument with an Agilent Eclipse Plus C18 column. A linear mobile phase gradient of 2-98% $CH_3CN$ (v/v) over 15 min in $H_2O$ with 0.1% formic acid (v/v) at a flow rate of 0.5 mL/min was used.

***Streptomyces tsukubensis* culture condition**: The seed culture of *S. tsukubenesis* was grown in TSB broth at 30 $^{o}$C for 24-36h. After that, the seed was transferred to 30 mL ISP 2 media with 1% volume ratio in a 150 mL Erlenmeyer flask and shaken for 5 days. After that, the culture was extracted with two equal amounts of ethyl acetate and dried. The dry extract was redissolved in methanol for HPLC-MS analysis with the same procedure as indicated in the above procedure.

**Quantitative analysis of isonitrile compound concentration**: Dry extracts (10 mL culture) of strain culture were redissolved in 400 μL methanol, to which 100 mM tetrazine in DCM was added with a final concentration of 10 mM. The mixture was stirred at room temperature for 3 h, and 5 μL of the final reaction solution was sampled and diluted 200 times in methanol for HPLC-MS (SingleQ).

***In-situ* reduction of isonitrile and tetrazine click reaction**: Dry extracts (2 mL culture) of strain culture were redissolved in 0.5 mL acetonitrile, to which 100 mM tetrazine in DCM was added with a final concentration 10 mM, and $NaBH_3CN$ in acetonitrile was quickly added to the above reaction with a final concentration of 30 mM. The reaction was allowed to stir for 0.5-1 h and then sampled for HPLC-MS (Q-TOF) analysis.

#### III. Quantitative analysis of universal product by HPLC-MS SingleQ

**1. Standard curve for universal product detection**: A series of pure universal product stock solutions (MeOH) with different concentrations (1mM, 3.125 mM, 6.25 mM, 12.5 mM, 25 mM) were prepared and subjected to HPLC-MS SingleQ analysis. Equivalent liquids were injected and the corresponding MS intensities were recorded. The mean value of triplicate runs was used to establish the standard curve.

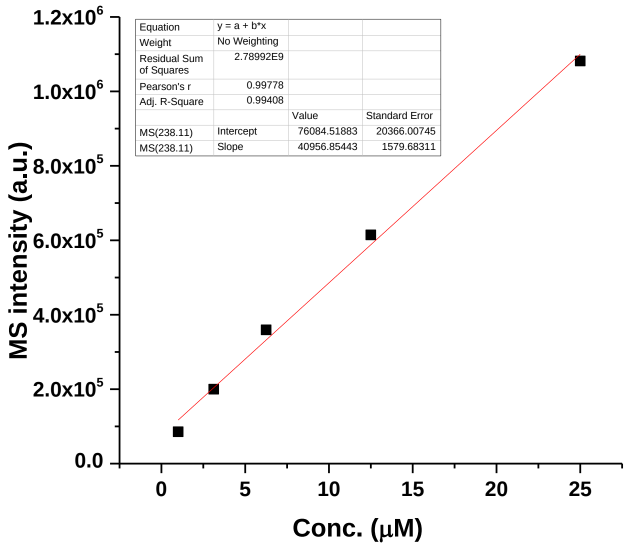


**Figure S1**. MS intensity-vs-concentration curve of the universal product.

**2. Rate constants calculation equations for the click reaction:** The isonitrile-tetrazine click reaction is a secondary reaction as revealed by previous papers [1].

$$\text{Secondary order reaction} \;\; v = k \times c[\text{A}] \times c[\text{B}]$$

$$\frac{1}{c} - \frac{1}{c_0} = k \times \text{t}$$

$$k = (\frac{1}{c_1} - \frac{1}{c_2})/(t_1 - t_2)$$

The sampling time were varied based on the actual reaction rates. For substrate **3** in MeOH solvent, k was calculated using the data recorded at 60 and 120 min of reaction time; For substrate **3** in DMSO or other mixed solvents, k was calculated using the data recorded at 20 and 30 min; For substrate **4** in MeOH or MeOH/$H_2O$, k was calculated using the data recorded at 0 and 15 min of reaction time.

**3. Color change of the click reaction**

Tetrazine shows a bright-red color while the universal product aminopyrazole shows a yellow color in common organic solvent (e.g. DCM or MeOH). When tetrazine was subjected to a reaction solution containing a certain amount of isonitrile (Py-tetrazine:isonitrile functionity >1:1

mole ratio), an obvious color change into light pink was occurred (mixed color of bright-red and yellow). If isonitrile compound was excessive, all tetrazine would be fully converted and resulted in a yellow color solution. (Cy-NC: cyclohexyl isocyanide)

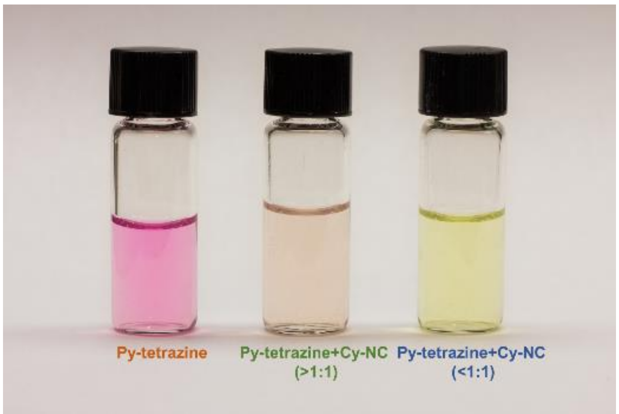


**Figure S2**. Color change of the isonitrile-tetrazine click reaction in methanol solution.

#### IV. Click reaction of INPLs from *E. coli-ScoA-E* and *E. coli-MmaA-E*

1. Click reaction with extracts from *E. coli-ScoA-E* culture

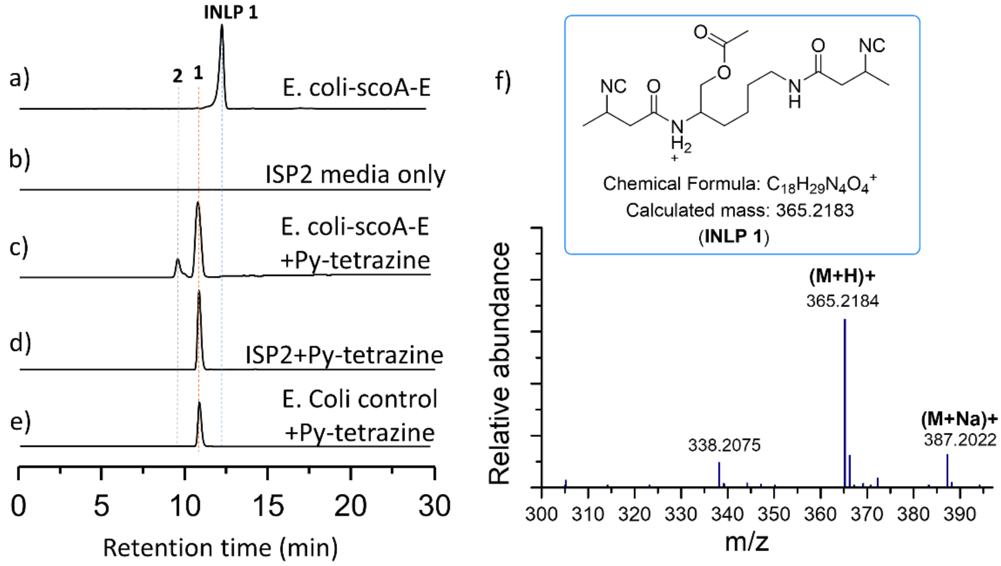


**Figure S3**. Click reaction for the identification of isonitrile compounds from *E. coli-ScoA-E* culture. Extracted ion chromatograms showing a) the production of **INLP 1** in the *E. coli-ScoA-E* culture extracts, b) no target compounds ISP2 media, c) full **INLP 1** conversion while the universal product **2** was detected, d) no reaction if Py-tetrazine **1** was added to ISP2 media only, and e) no reaction when the extracts of E. coli control strain that does not produce isonitrile metabolites was used to react with Py-tetrazine **1**. The calculated masses for **INLP 1** (365.2183), Py-tetrazine (237.0883) and universal product (238.1087) with 20 ppm mass error tolerance were used for each trace. f) structure and MS results of the **INLP 1** product.

2. Click reaction with extracts from *E. coli-MmaA-E* culture

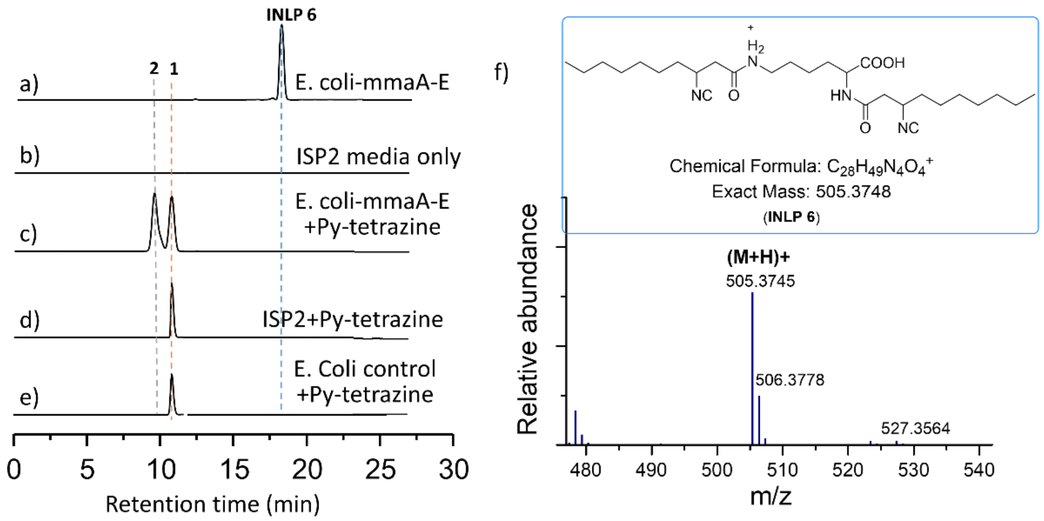


**Figure S4**. Click reaction for the identification of isonitrile compounds from *E. coli-MmaA-E* culture. Extracted ion chromatograms showing a) the production of **INLP 6** in the *E. coli-MmaA-E* culture extracts, b) no target compounds in ISP2 media, c) full **INLP 6** conversion while the universal product **2** was detected, d) no reaction if Py-tetrazine **1** was added to ISP2 media only, and e) no reaction when the extracts of E. coli control strain that does not produce isonitrile metabolites was used to react with Py-tetrazine **1**. The calculated masses for **INLP 6** (505.3748), Py-tetrazine (237.0883) and universal product (238.1087) with 20 ppm mass error tolerance were used for each trace. f) structure and MS results of the **INLP 6** product. f) structure and MS results of the **INLP 6** product.

#### V. Titer quantification of isonitrile compounds from strain culture extracts.

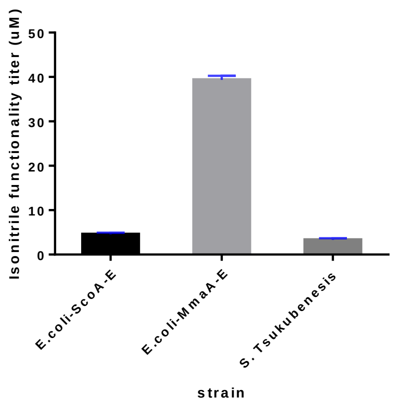


**Figure S5.** Titers of the isonitrile compounds in different stain cultures.

#### VI. Identification of isonitrile compounds from *Streptomyces tsukubenesis NRRL 18488* strains.

1. Culture media screening through click reaction.

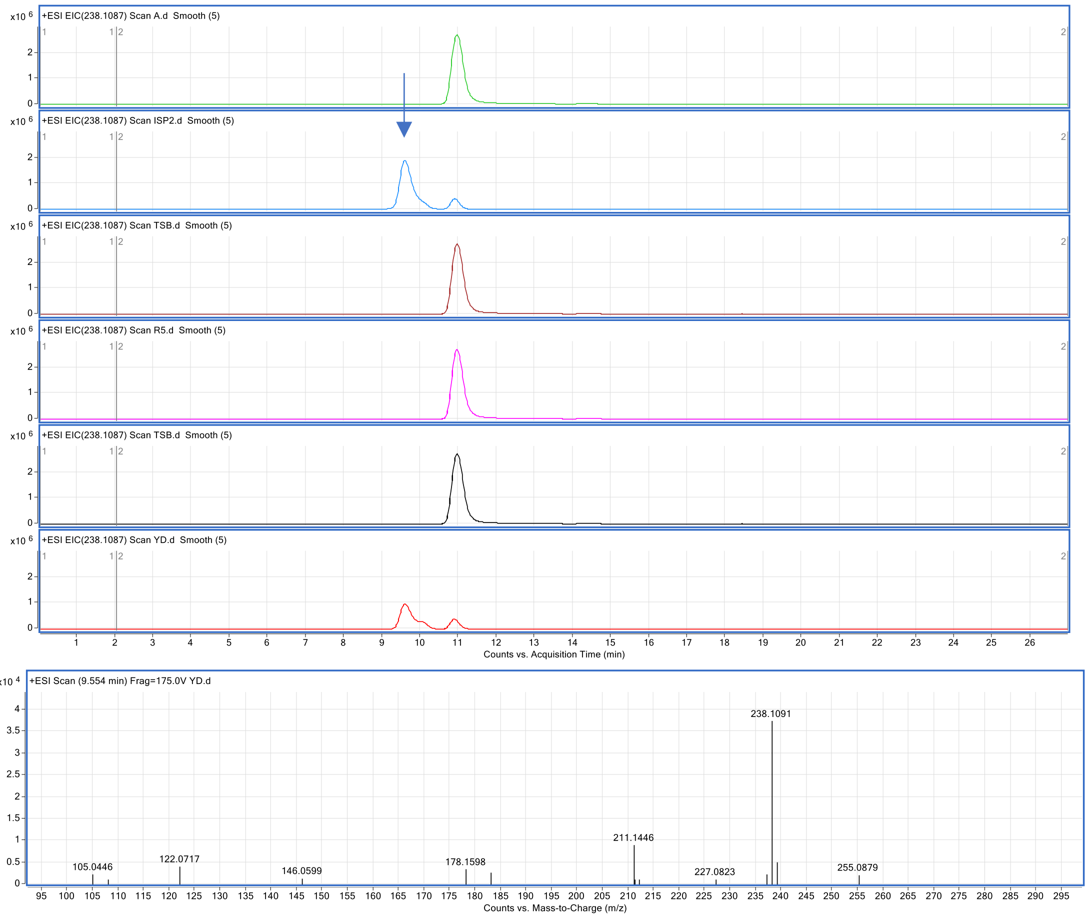


**Figure S6**. Click reaction of the *S. tsukubenesis* culture extracts with different media. From top to bottom: A media, IPS2 media, R5 media, TSB media, YD media. Two (ISP2/YD) out of six media benefit the production of universal product after click reaction. EIC trace of the molecule weight 238.1087 (universal product).

2. MS analysis of the isonitrile compound

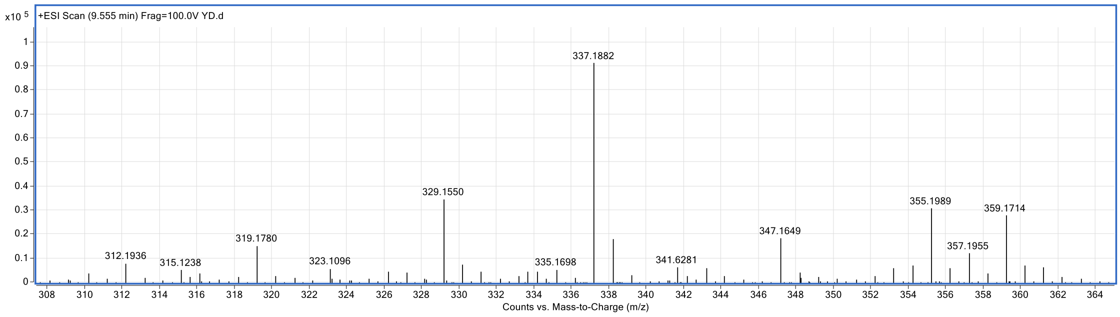


**Figure S7**. QTOF mass analysis of the isonitrile compound SF2768 in ISP2/YD media.

#### VII. *In-situ* reduction of the click reaction

1. *In-situ* reduction of the click reaction with *E. coli-MmaA-E* extracts.

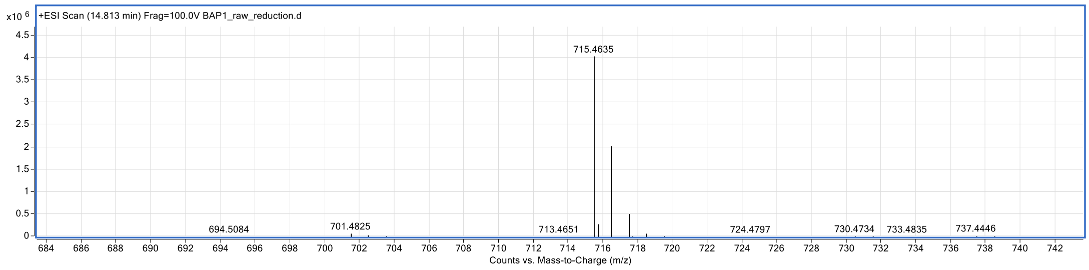


**Figure S8.** HRMS analysis of the mono-conjugate **6** from the *in-situ* reduction of the click reaction between *E. coli-MmaA-E* and Py-tetrazine (Calculated 715.4654, found 715.4635).

2. *In-situ* reduction of the click reaction with *E. coli-ScoA-E* extracts.

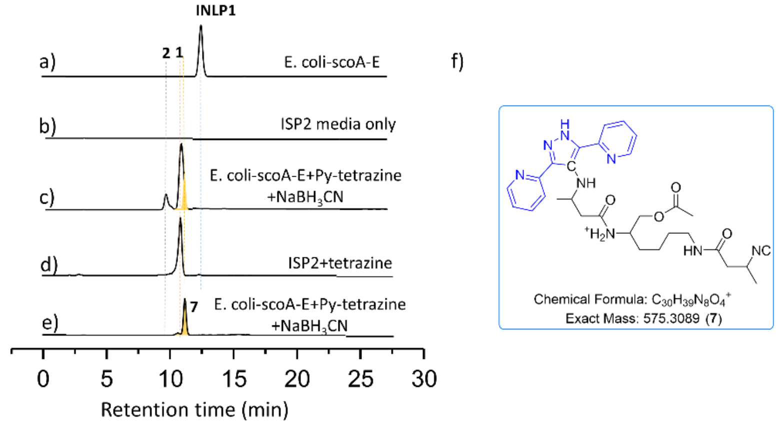


g) QTOF data:

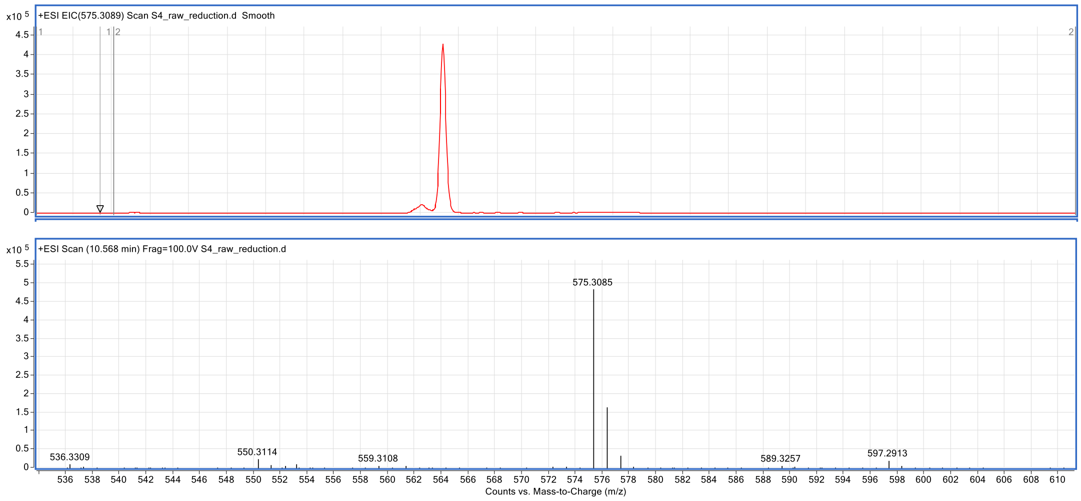


**Figure S9**. *In-situ* reduction of the click reaction between *E. coli-ScoA-E* culture extracts and Py-tetrazine. Extracted ion chromatograms showing a) the production of **INLP 1** in the *E. coli-ScoA-E* culture extracts, b) no target compounds in ISP2 media, c) full **INLP 1** conversion while the universal product **2** and mono-conjugate **7** was detected, d) no reaction if Py-tetrazine **1**

was added to ISP2 media only. The calculated masses for **INLP 1** (365.2183), Py-tetrazine (237.0883), universal product (238.1087) and mono-conjugate **7** (575.3089) with 20 ppm mass error tolerance were used for each trace. e) Extracted ion chromatograms showing the presence of mono-conjugate **7.** The calculated masses for mono-conjugate **7** (575.3089) with 20 ppm mass error tolerance was used for this trace. f-g) predicted molecule structure and MS results of mono-conjugate **7**.

3. *In-situ* reduction of the click reaction with *S. tsukubenesis* extracts.

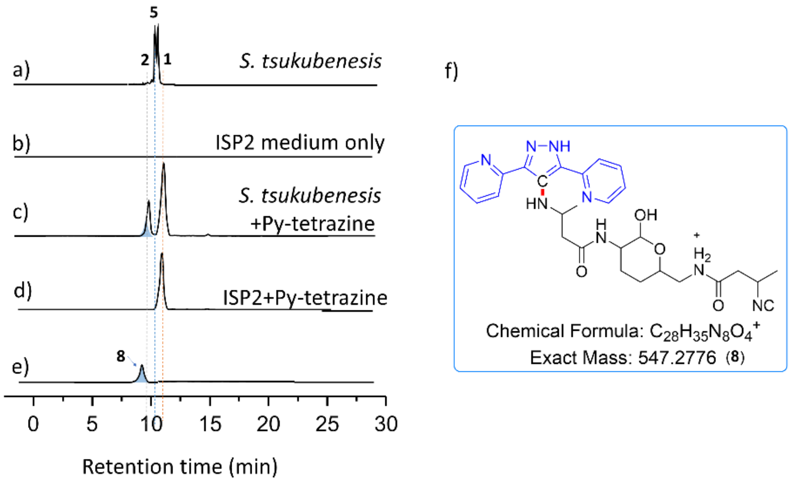


**g)** QTOF data

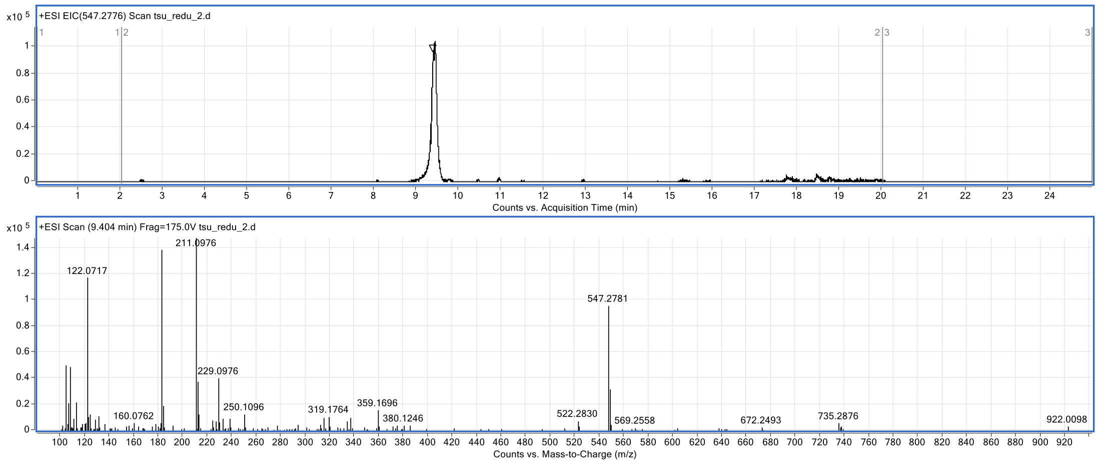


**Figure S10**. *In-situ* reduction of the click reaction between *S. tsukubenesis* culture extracts and Py-tetrazine. Extracted ion chromatograms showing a) the production of **SF2768** (**5)** in the *S. tsukubenesis* culture extracts, b) no target compounds in ISP2 media, c) full SF2768 conversion while the universal product **2** and mono-conjugate **8** was detected, d) no reaction if Py-tetrazine **1** was added to ISP2 media only. The calculated masses for **SF2768** (337.1870), Py-tetrazine (237.0883), universal product (238.1087) and mono-conjugate **8** (547.2776) with 20 ppm mass error tolerance were used for each trace. e) Extracted ion chromatograms showing the presence of mono-conjugate **8.** The calculated masses for mono-conjugate **8** (547.2776) with 20 ppm mass error tolerance was used for this trace. f-g) predicted molecule structure and MS results of mono-conjugate **8**.

H-NMR characterization of the universal product

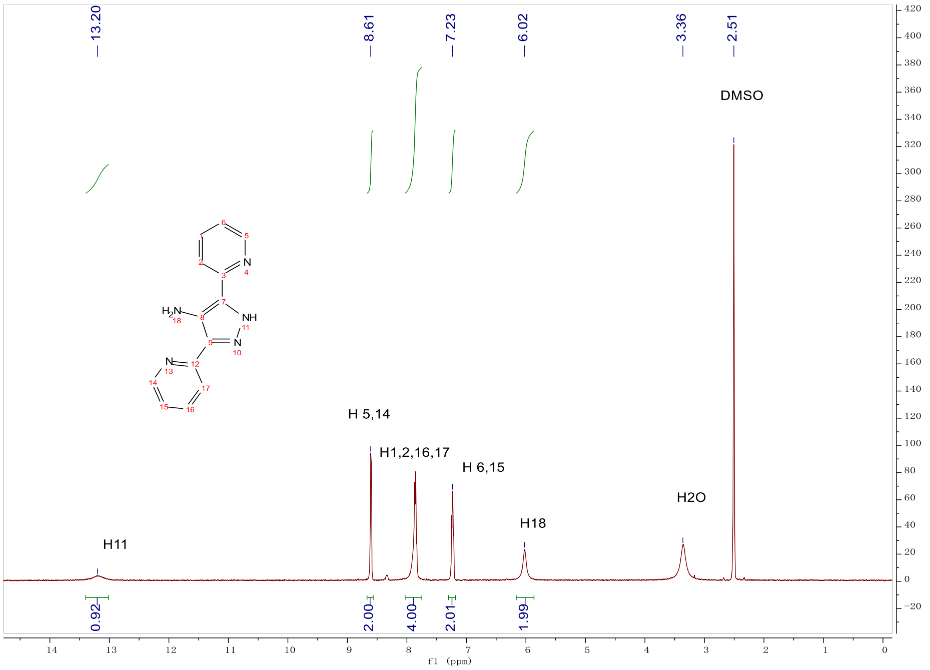
